# Molecular Dynamics Simulations Reveal the Importance of Non-Native Interactions in Modulating the Inactive to Active Conformational Transition in Progesterone Receptor

**DOI:** 10.1101/2024.10.18.619098

**Authors:** Saurov Hazarika, C. Denise Okafor

## Abstract

Nuclear receptors are a family of transcription factors that activate respective genes when bound to specific ligands. Certain ligands induce conformational changes in the receptor which helps them recruit co-activators. The conformational changes induced by these ligands conformationally transition inactive conformation into an active conformation by changing the orientation of the C-terminal helix (H12). Despite their immense physiological importance, very few questions have been solved about the kinetics and the molecular mechanism of this transition from the inactive to active conformation. In this study, we have used extensive unbiased atomistic molecular dynamics simulations of Progesterone receptor bound to a partial agonist asoprisnil to investigate these two questions. Two different crystal structures for this complex provide us with a unique opportunity to study the conformational transition at the molecular level. Apart from elucidating several important dynamical information from these simulations, we used Markov state modeling to calculate the rate of the transition between the inactive and active-like states. More importantly, we have also shown the importance of non-native interactions in this conformational transition, which were seen to be formed during the transition from inactive to active-like conformation but not present in the active conformation itself. Apart from contributing to our fundamental understanding about the structure and dynamics of nuclear receptor at the molecular level, this study might be able to contribute to the larger problem of protein-folding itself.

## Introduction

Nuclear receptors are secondary transcription factors that transactivate genes when bound to specific ligands. When specific ligands bind to these nuclear receptors, they induce conformational changes in the receptor and the nuclear receptors get activated. Nuclear receptor has a multi-domain structure, consisting of structured regions such as Ligand-binding domain and DNA binding domain and the unstructured regions such as the hinge domain and the N-terminal domain. Ligand binds to a hydrophobic ligand binding pocket which is situated at the ligand binding domain of the receptor. It was earlier shown that the active and inactive receptor can be recognized based on their conformation. According to the classical activation model of nuclear receptor, the c-terminal helix or the helix 12 (H12) of the receptor is differently oriented in active and inactive conformation^1–4^. Helix 12, helix 3 and helix 4 form the activation surface 2 where the co-activator binds. In the active conformation, H12 is docked against the helix 3 and helix 4, that together forms a conducive surface for the co-activator to bind. On the other hand, in the inactive conformation, helix 12 shifts away from helix 3 and helix 4, making it difficult for co-activator to bind while facilitating the co-repressor binding.

Using experimental techniques such as NMR and HDX-MS, previously it was shown that the H12 region is conformationally fluid. Different orientations of H12 have been observed in crystallographic studies. Several crystal structures have been shown to contain an extended H12 region where H12 is completely shifted away from the rest of the ligand binding domain ^1,5^. The fact that it was first observed in the apo structure prompted some to assume the extended conformation as the “unliganded” conformation. But later on, the extended structure has been observed in holo nuclear receptor as well^1,6^. It also raises doubt over the physiological relevance of the extended conformation. Later, several apo structures were crystalized in the active conformational state as well. It makes it more difficult to predict how H12 orientation is modulated by the binding of different ligands. In a broad sense, the extended state where H12 regions is shifted away from H3 and H4 region is considered as “inactive” conformation and the conformation where H12 is docked against H3 and H4 region is considered as “active” conformation^5,7,8^.

Ligands that activate the nuclear receptor by helping the receptor to recruit co-activator are known as “agonist” and the ligands that deactivate the receptor by blocking the recruitment of the co-activator are known as “antagonist”. Then there are ligands that don’t fall into the classical categories of “agonist” or “antagonist”. This category includes partial agonists, mixed agonists/antagonists and selective modulators. Based on different parameters, these ligands have the ability to both activate and deactivate a particular receptor; thus, they sit somewhere between “agonist” and “antagonists”. Such examples were seen for PPAR_γ_ and VDR^9–11^.Progesterone receptor was also crystalized in both active^12^ and inactive conformation^13^ when bound to the ligand Asoprisnil, which is a selective PR modulator. In the PR-asoprisnil antagonist conformation, when a co-repressor peptide is bound, H12 regions assumes a unique position where although extended, it is docked against the H11 region. Having crystal structures for both active (PDB 4A2J) and inactive conformation (PDB 2OVM) for the single ligand provides a unique opportunity to study the dynamics of conformational transitions between active and inactive conformation. Taking advantage of this specific point, in an earlier study, we used accelerated molecular dynamics simulations and enhanced sampling method to study the dynamics and the energetics of this conformational transition in detail^14^. Using accelerated molecular dynamics simulations, we have shown that the transition from inactive to active conformation happens at a long-time scale. Furthermore, using umbrella sampling, we have shown that the active conformation represents the global minima in the free energy surface along a certain reaction co-ordinate while the inactive conformation represents one of the metastable states. But a detailed kinetic study is still lacking since it’s not possible to calculate the rate of the transition using a biased simulation and PMF is not enough to make a comment about the exact kinetics of the system. Moreover, the molecular reason behind this difference in stability between these two conformations is yet to be answered. Using a large set of classical atomistic simulations, we have tried to answer them in this study.

In this paper, we have first used several unbiased atomistic MD trajectories to study the conformational transition between the active and inactive conformations. A small set of unbiased MD trajectories is statistically inadequate to study this transition. Initial studies have recapitulated our findings from the accelerated molecular dynamics simulations. A large set of MD trajectories help us in constructing the Markov state model to calculate the kinetics of the transition between the active and inactive states. The Markov state model is a statistically rigorous framework to study the kinetics of the conformational transition in MD simulations ^15^. Using Markov state model, we were able to show rigorously that this transition happens at the microsecond scale.

The other question that we have addressed is the molecular forces that drive the transition from inactive to active-like conformations. The inter-residue interactions such as salt-bridges and hydrogen bonds formed by the residues present in H12 and L11-12 regions seem to play a significant role in making such transition possible. We were able to show that a set of non-native interactions modulate this conformational transition. In this case, we consider the active conformation as the native conformation. At the molecular level, we have seen the formation of non-bonded interactions such as salt-bridges and hydrogen bonding during the process of conformational transition, but these interactions are not present during the simulations of the active or native conformation itself. Whether non-native contacts or interactions play a role in protein folding is a matter of big debate^16–19^. Although our current study isn’t done to study the protein folding in general term, having two crystal structures of the same protein complex provide us with a unique opportunity to study the conformational transition into native structure from the non-native one at the molecular level, which in turn may turn out to be an important contribution in the problem of protein folding as well. Based on our findings, we have come up with a toy thermodynamic model which might explain the process of inactive to active conformational transition in Progesterone receptor.

## Methods

### Structure preparation

The Crystal structures of PR bound to partial agonist were taken from the RCSB PDB. The PDB ID of the active conformation is 2ovm and the PDB ID of the inactive conformation is 4a2j. To add any missing residues, we used Modeller^20^. In our simulations and subsequent analysis, we renumbered the PR LBD residues 683-932 to 1-250 for simplicity. Simulations were performed on these two complexes.

### Classical MD simulations

The ligand, Asoprisnil, was parameterized using Antechamber tool^21^ and Generalized Amber Force Field 2 (GAFF2)^22^ available in AMBER22. The complex is solvated using TIP3P water model with 10 Å buffer. To attain the physiological salt concentration of 150 mM, Na^+^ and Cl^-^ ions were added to the system. Four steps of minimization were performed to the solvated complex. In the first step, 500-kcal/mol. A^2^ restraining force was applied to the solute atoms (Protein and the ligand) during the minimization process. In the second step, the restrain was reduced to 100-kcal/mol. A^2^. In the third step, the restrain was removed from the protein and a 100-kcal/mol. A^2^ restrain was applied to the ligand only. In the fourth step, all restraints were removed from the system. To do the minimization, 5000 steps of gradient descent followed by 5000 steps of conjugate gradient minimization were applied in each step. After minimization, the system was heated from 0 to 300K in a 100 ps of NVT simulations. A 5-kcal/mol. A^2^ restrain force was applied to all protein and ligand atoms during the heating. The NVT simulations were performed in three sequential steps. In the first step, 10 kcal/mol. A^2^ restrain was added to protein and ligand. In the second step the restraint was reduced to 1 kcal/mol. A^2^. In the third step, the restrain was limited to only ligand atom. After this the restraints were removed. 500ns to 1 microsecond production run were performed using a 2-fs time step. SHAKE algorithm^23^ was applied as a distance constraint to all the bonds between heavy atoms and hydrogen atoms. For all the simulation steps, ff14SB force field was used^24^. The simulations were performed in AMBER22 software ^25,26^. Different python modules such as MDtraj^27^, MDAnalysis, Pyemma^28^ were applied to calculate RMSD, RMSF, DSSP, contact maps, PCA and -log(probability) plots.

### Markov state modelling

Pyemma^28^ python module was used for tICA and MSM analysis. K-means clustering was chosen to discretize the sample space. Three features have been selected for our MSM-the minimum RMSD value with respect to the inactive conformation, the minimum RMSD value with respect to the active conformation and the minimum distance between two specific segments of the protein which shown to come closer in the active conformation while stays apart in the inactive conformation. The further statistical tests required for building a MSM are in supplementary document submitted with this paper (Fig S1).

## Results

To monitor the conformational changes in active and inactive conformations, we obtained 50 500ns classical MD trajectories of both active and inactive PR-Asoprisnil complexes, allowing us to achieve statististical rigor. Fig -1(A) and 1(B) are the inactive and active conformations of the receptor-ligand complex. The RMSF analysis identified L11-12 region as the most flexible region (Fig 1E). Among the structured regions, H12 helix region shows the highest flexibility, recapitulating earlier results of accelerated MD simulations^14^. It can also be seen that for the inactive complex, H12 and L11-12 regions have much higher fluctuation compared to the active complex (Fig 1E) while other regions have almost similar fluctuations. To further analyze the structural stability of the H12 region, DSSP analysis was perfomed for both complexes. The H12 region of the active complex shows a higher tendency in forming helical structure compared to the inactive complex (Fig S3) indicating higher structural stability of the H12 region in the active complex.

**Fig 1:**
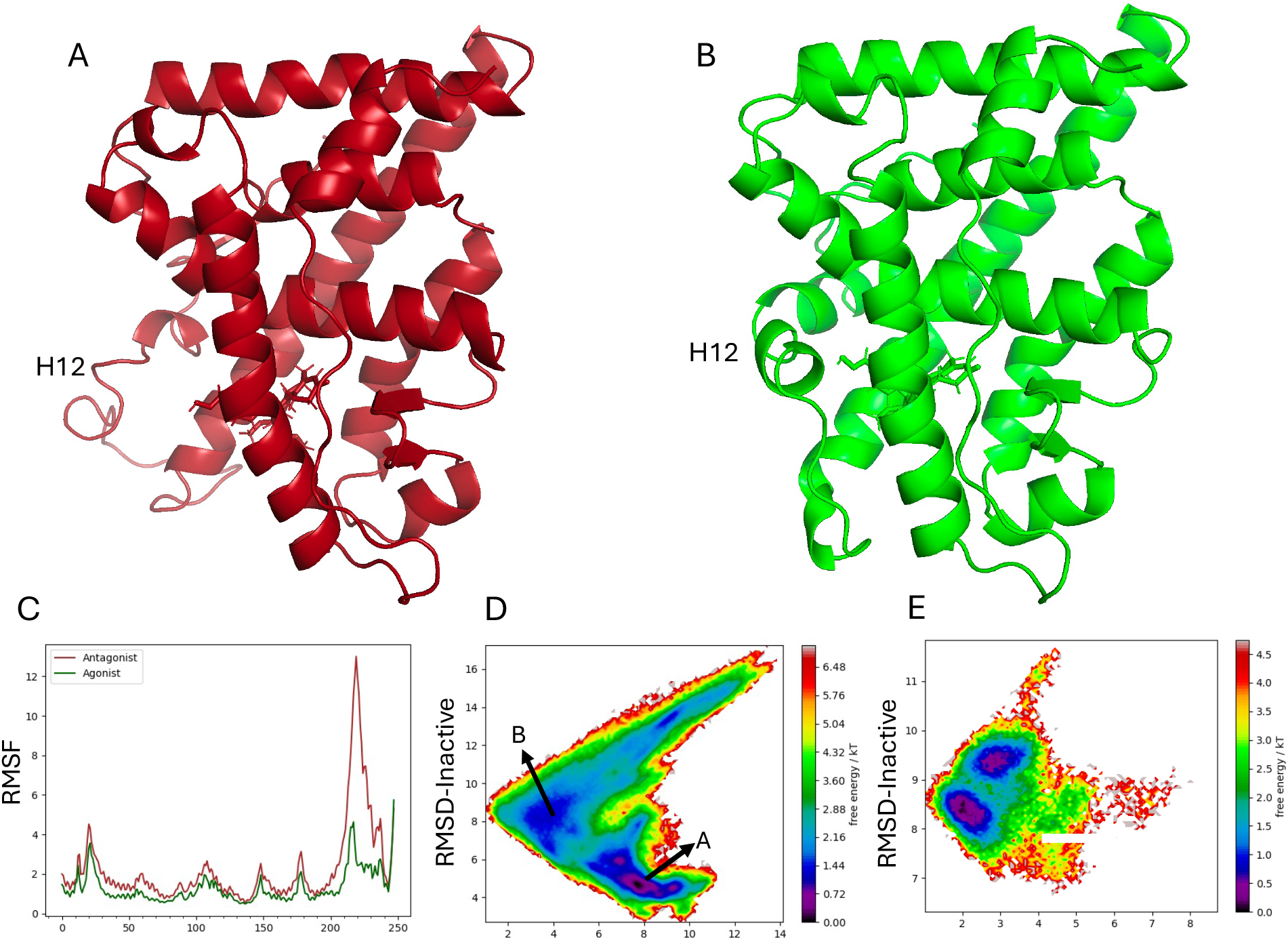
Classical Molecular Dynamics simulations of active and inactive conformations of PR-Asoprisnil complex. (A) and (B) PR-asoprisnil inactive and active conformations. The orientation of the H12 and L11-12 regions mainly differentiates these two conformations. (C) RMSF plot for active and inactive conformation. H-12 and L11-12 region show the highest flexibility for both complexes. Inactive complex shows higher flexibility in those regions compared to the active complex. (D) and (E) -log(probability) plots for both inactive and active conformations when bound to asoprisnil. Y-axis stands for the RMSD of the H-12 region with respect to the inactive conformation while the x-axis stands for the RMSD of the H-12 region with respect to the active conformation.

To characterize the structural transitions between the two structures, RMSD analysis was performed. For both the complexes, RMSD was calculated with respect to both the initial active and inactive structures. For example, for the trajectories of the active structure, the two types of RMSD were calculated. For the first one the reference structure is the initial active conformation while for the second one the reference structure is the initial inactive conformation. -Log (Probability) shows two energy minima for the inactive complex (Fig 1D). In the first minima (A), RMSD-inactive is lower than RMSD-active, suggesting a conformation that is closer to the inactive complex. In the second minima (B), the reverse situation is observed, identifying the transition of the inactive complex to an active-like state over time. For the active complex, although the -log (Probability) plot shows two minima, both cases show RMSD-active as lower than RMSD-inactive (Fig 1E). Thus, the active conformation has a higher likelihood of remaining in this state while the inactive complex tends to transition into the active conformation. This finding suggests a greater stability in the active conformation.

A dynamic cross-correlation analysis shows multiple regions where the active complex has weaker correlation compared to the inactive complex. (Fig 2A and 2B). For example, L8-9 loop region is dynamically correlated with the H3 and L3-4 loop regions in both cases. Similarly, the H1 region is dynamically correlated with H8, H9 and L8-9 regions. We also see a weak correlation of H8, H9 and L8-9 regions with part of the H12 region.

**Fig 2:**
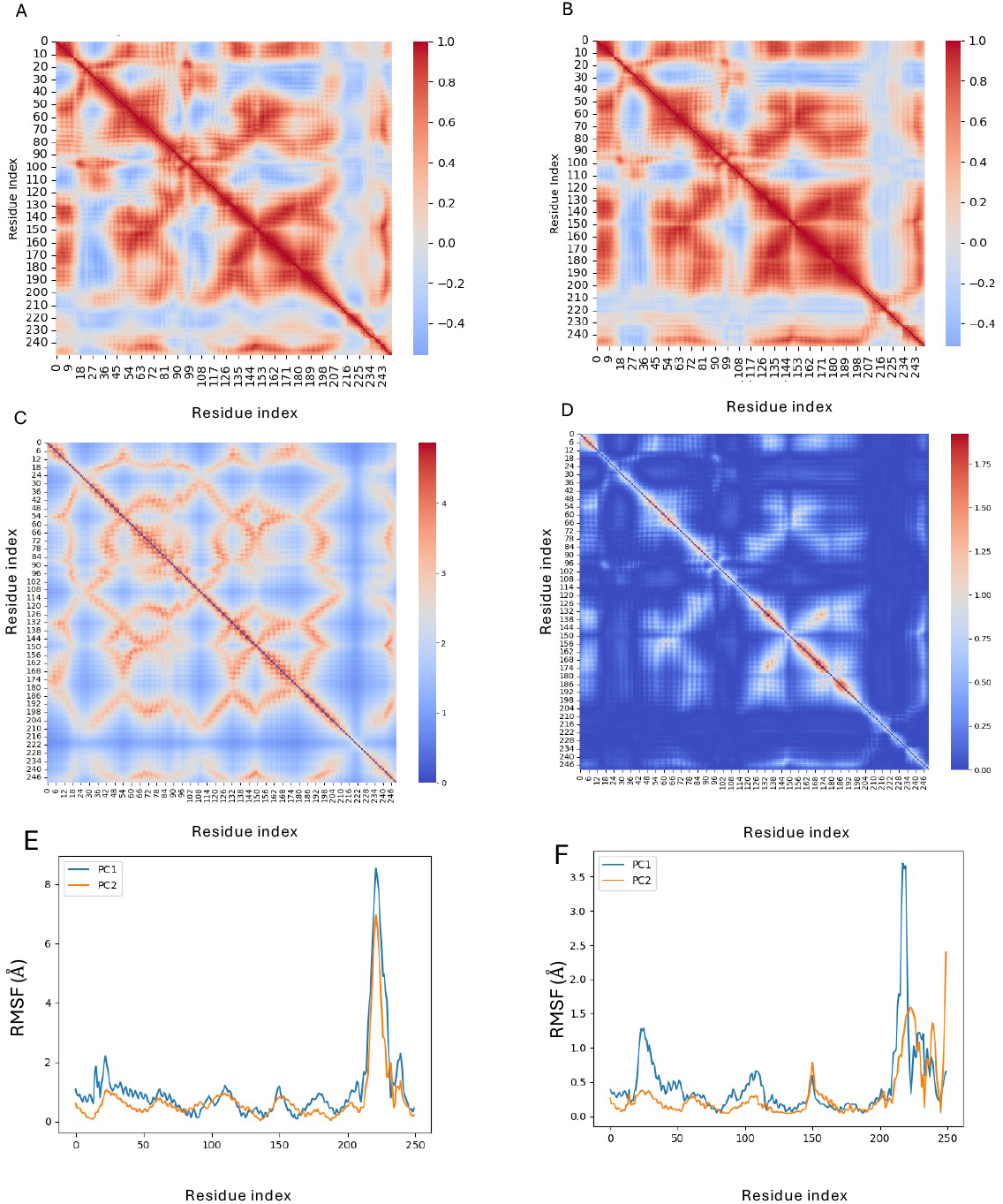
(A) and (B) Dynamic cross-correlation analysis of inactive and active conformations respectively. (C) and (D) Linear Mutual Information (LMI) based cross-correlation analysis of the inactive and active conformation respectively. (E) and (F) The RMSF plots of the Inactive and active conformations respectively along PC1 and PC2.

To incorporate the correlation in the orthogonal direction, which is ignored by dynamic cross-correlation analysis, we have done linear mutual information (LMI) based cross-correlation analysis as well (Fig 2C and 2D). The reason behind doing these two analyses is to understand how different regions in the protein are correlated and how these correlated regions change for active and inactive conformations. Although the dynamic cross-correlation analysis doesn’t show a stark difference in correlation between different regions of inactive and active conformations, the LMI correlation is clearly much larger for the inactive conformation compared the active conformation. H12 region in the inactive conformation is strongly correlated with H10/11, L8-9 and H5 regions. Other such strong correlations can be observed between L4-5 and H8/9 regions, H3 and H5/6 regions, and H1 and H9 regions. In the case of active conformation, moderate correlation can be observed between H7/8 and H9/10/11 regions, H4/5 and H9 regions, and H1 and H9 regions. Moderate correlation was observed between H12 and H8/9/10 regions as well. The correlation of the H12 region with other parts of the protein is much higher in the case of inactive conformation compared to the active conformation.

To better understand the high dimensional simulation data, principal component analysis (PCA) was performed for both complexes. The motions described by PC1 and PC2 differ widely between the two complexes. In the case of the inactive complex though, H12 and L11-12 regions have high fluctuations along both PC1 and PC2 (Fig 2E). For the active complex, while the overall fluctuation of H12 and L11-12 regions are less than the inactive complex, the fluctuation of the L11-12 region is higher along the PC1 than the PC2 (Fig 2F).

### Markov state modelling shows that the transition from inactive to active conformation happens in microsecond time scale

In our previous report, we established the active conformation as the global energy minima along a particular reaction-coordinate and also used 500-ns accelerated MD simulations to describe the transition from inactive to active states ^14^. While that study hinted at a transition timescale on the order of microseconds, it is not possible to calculate a more exact timescale using accelerated MD simulations. To study the inactive to active (and vice-versa) conformational transition of the protein of interest in more detail, Time-lagged independent component analysis (TICA) was applied to our unbiased MD simulations. The goal of this operation was to find a set of features that best describe this conformational transition and based on which other factors at the molecular level can be determined. The simulations data were projected in a feature space consisting of the minimum distance between the L11-12 and L2-3 loops and the minimum RMSD value with respect to the active conformation. The TICA components were used to cluster the simulations data into a large number of microstates, which were then used to construct the MSM.

Markov state modelling provides a statistically robust way to calculate the kinetics in molecular dynamics simulations^15^ by using multiple trajectories for better sampling and statistical rigor. In this study, we build Markov models for both complexes using 50 trajectories (totaling 25 microseconds) of each complex.

First, we focus on the inactive complex, with statistical tests suggesting the presence of two distinct states (S1 A and B). The tICA projection plots (Fig 3A) along IC1 and IC2 show two distinct states for the inactive complex. Representative structures identify the states as near-active and inactive conformations, respectively (Fig 3B-C), where the inactive conformation occupies a significantly larger area of the tICA plot compared to the active state. We note that while constructing the MSM, all simulations were initiated from the same inactive structure. Thus, the trajectories spend more time sampling the initial conformation than any other regions of conformational space, explaining the larger proportion of the inactive conformation in the tICA plot. The transition time from the inactive state to the active state is calculated as 9072.4 ns or 9.07 microseconds (Fig 3D), confirming that the transition happens at the microsecond time scale.

**Fig 3:**
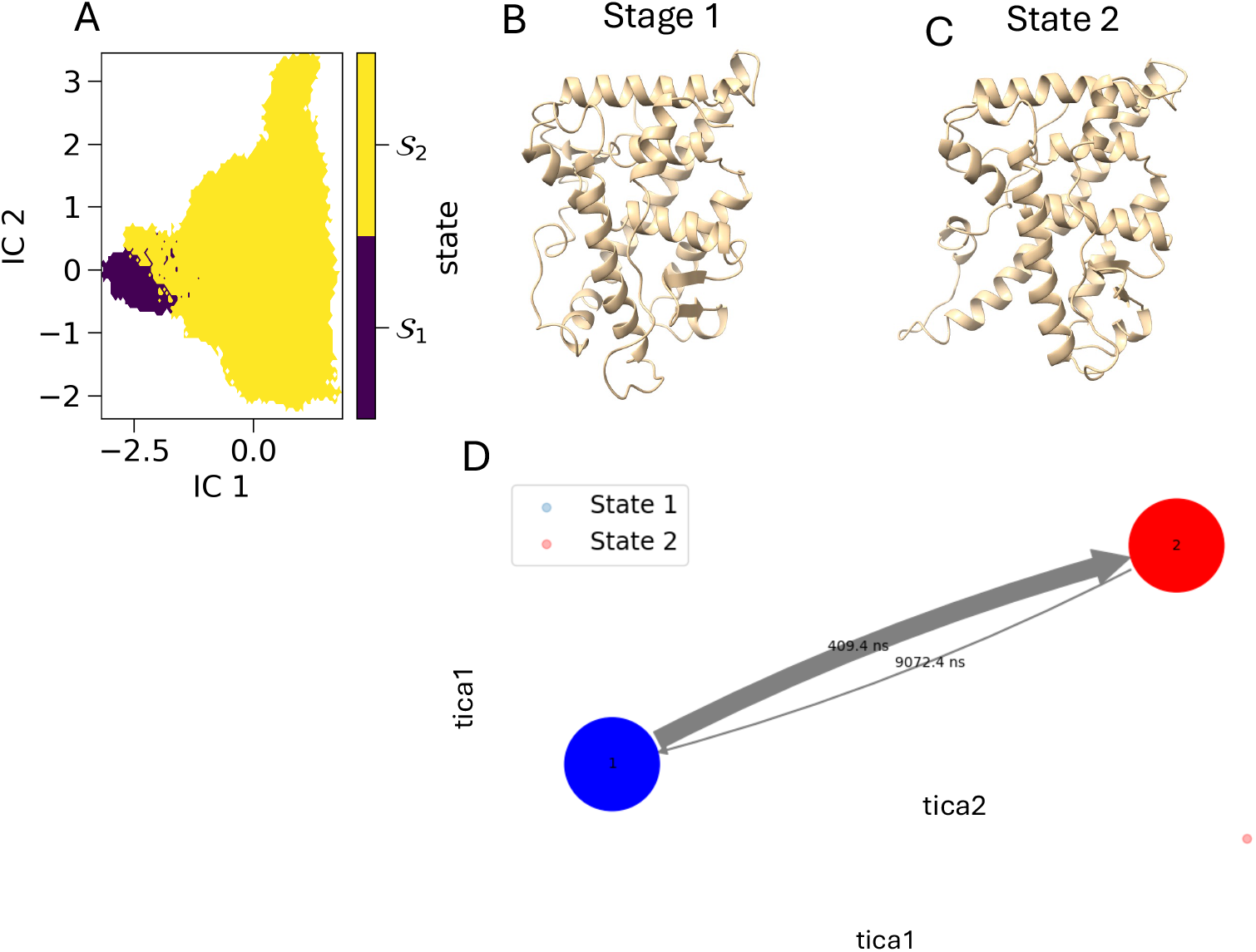
Markov state modelling of the inactive complex (2ovm). (A) PCCA distribution of tICA projection. The plot shows two well-separated states. (B) and (C) Representative structures of State 1 and State 2 generated in MSM. State 1 resembles the active conformation while state 2 resembles the inactive conformation. (D) Transition between states generated in MSM.

Despite running similar number trajectories, we didn’t see a transition from active to inactive conformation when started from the active complex. Two states in the tICA plot for the active complex contain conformations that are very much like the active conformation (Fig S2). This recapitulates the earlier finding that the active conformation being the global minima in the configurational space, have difficulty in transition into a higher energy inactive state and much longer simulation time is needed for this transition.

### Non-native slat-bridge formation may modulate the inactive to active conformational transition

Next, we aim to study the conformational transition observed for the inactive conformation at the molecular level and try to dissect any possible molecular forces that modulate this transition. Considering the fact that the active conformation is more stable than the inactive conformation, we’ll assume that the active conformation represents the native structure of the protein complex in this case. To check whether our conclusion from the previous section that the state 2 in the TICA plot really represent a near-active conformation or not, we have calculated the fraction of native contacts (*Q*) with considering active conformation as our native state and projected this value into TICA plot. The fraction of native contacts tells us how close our current structure is to the native structure, and we used the formula derived in^19^ to calculate the fraction of native contacts. In our all-atom model, a contact is formed when the distance between any pair of heavy atoms *i* and *j* of residues *θ*_*i*_ and *θ*_*j*_ is less 4.5 Å provided |*θ*_*i*_ -*θ*_*j*_| > 3. Then Q is defined as^19^:

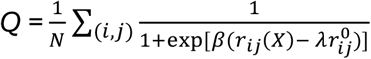

Q was calculated for each frame in every trajectory of the Inactive conformation and the values were projected into the TICA plot. It’s clear from the Fig-4 that as we move from state 1 to state 2, the fraction of native contacts increases and thus the conformation is becoming more native like.

**Fig 4:**
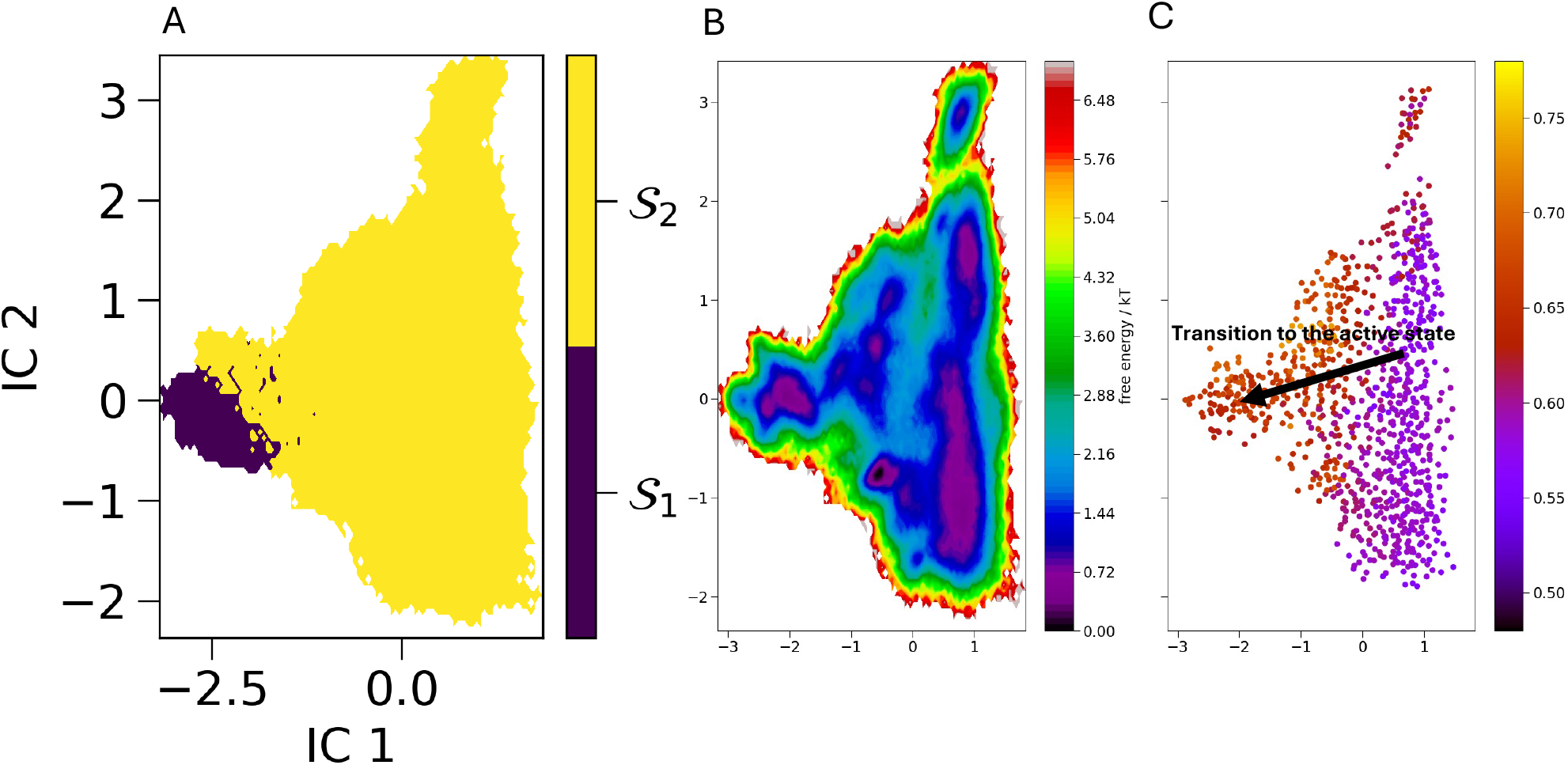
The fraction of native contacts projected into the energy landscape along first and second time-lagged independent components. (A) is the PCCA distribution of the TICA projection to compare visually which part of the energy landscape represents the state 2 and which part represents the state 1. (B) the energy landscape and (C) the fraction of native contacts projected into the energy landscape. The fraction of native contacts increases while going from state 1 to state 2, which represents an inactive to active conformational transition while moving from state 2 to state 1.

Since the highest fluctuations were observed in H12 and L11-12 regions, we tried to check if there was any type of outstanding non-bonded interactions present in the region apart from hydrophobic interactions, that may play a role in modulating the inactive to active conformational transition. We randomly selected ten representative structures from the two states (state 1 and state 2) and used ProteinTools^29^ to visually inspect the salt-bridges formed in those structures. The goal here was to detect any salt-bridges that have a dominant presence in state but absent in the other state. We observed two such salt-bridges formed by the residues of L11-12 region.

Salt-bridges are electrostatic interactions formed between negatively charged sidechain of the acidic residues (Asp and Glu) and the positively charged sidechain of the basic residues (Lys and Arg). Specifically, we observed the prominent presence of the GLU 222 – ARG 42 salt-bridge where GLU 222 resides in the L11-12 loop and ARG 42 resides in H3 region. Another salt-bridge that we witnessed is ARG 217 – GLU 41, where ARG 217 resides in L11-12 region and GLU 41 resides in the H3 region. The first salt-bridge, i.e. GLU 222 – ARG 42, is very much absent in the inactive conformation but has a dominant presence in the near-active conformation. The other salt-bridge, i.e. ARG 217 – GLU 41, could be seen forming in both states but its presence in state 2, i.e. the near active conformation is much higher than that of the inactive conformation. To quantify the number of salt-bridges and to check if there is any difference in its presence in the two states, we calculated the number of salt-bridges formed between these residues. We have used a distance cut-off of 4.5 Å between oxygen atoms of the acidic sidechain and nitrogen atoms of the basic sidechain to define a salt-bridge. Although in the stability of the salt-bridge the angle might play a minor role by providing an extra strength in terms of hydrogen bond, in our case we used the distance as the sole criteria to define our salt-bridge. Once we separately calculated the number of these two salt-bridges formed during our simulations for each trajectory, we projected them into our TICA plot. The difference of the presence of these two salt-bridges in two states is apparent from the plot. For GLU 222 – ARG 42 salt-bridge, its presence can be seen only in the near-active region (Fig 5 A, B and C). On the other hand, the ARG 217 – GLU 41 slat-bridge can be seen forming in both states but the inactive state its formation is very unstable (Fig 5 D, E and F). The black dots in the plot represent the conformations with zero salt-bridges. While in state 1 or in the inactive conformation the formation and disruption of this salt-bridge is more common, in state 2 or the near-native conformation ARG 217 – GLU 41 salt-bridge is very stable and has a prominent presence. This establishes the importance of these interactions in transitioning from the inactive to active conformation. To further analyze the importance salt-bridges formed by H12 and L11-12 residues, we selected a trajectory from the set of fifty trajectories of the inactive complex that shows the transition from inactive to active conformation and calculated the salt-bride formation in every frame. From the figure (Fig S4) number of interactions sharply increases with time, i.e. while going from the inactive to active-like conformation.

**Fig 5:**
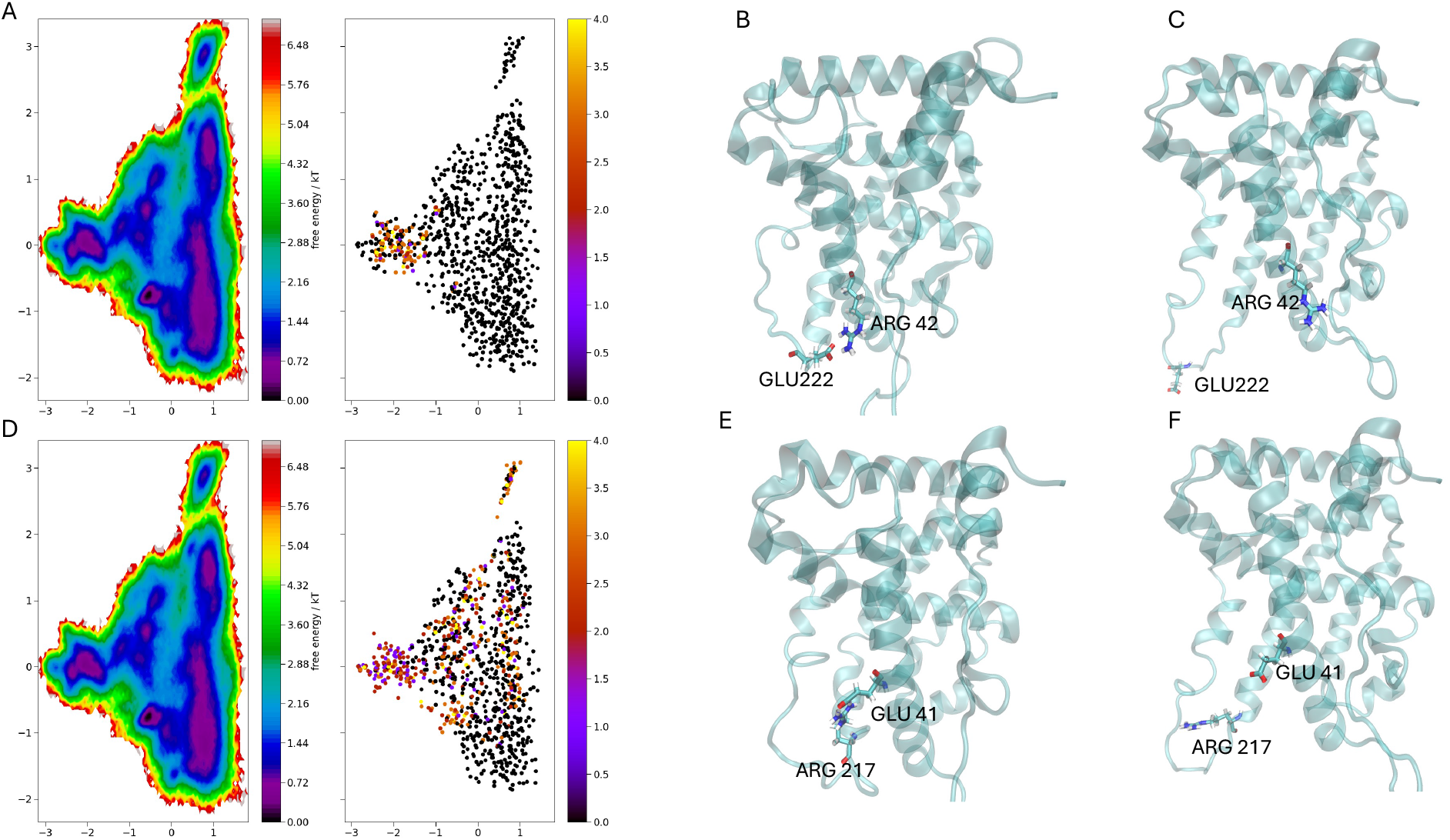
Salt-bridge formation in different protein conformations. (A) GLU 222 – ARG 42 salt-bridge formation projected into the energy landscape. The number represents all the possible contacts where the oxygen atoms of the acidic sidechain and nitrogen atoms of the basic sidechain are within 4.5 Å. The number increases while going from the state 2, i.e. the inactive conformation, to state 1, i.e. active-like conformation. (B) A representative structure from the state 1, i.e. active-like conformation, where GLU 222 and ARG 42 can be seen close to each other, forming a salt-bridge. (C) A representative structure from state 2, i.e. the inactive conformation, where GLU 222 and ARG 42 can be seen situated far apart from each other, and thus unable to form any contact. (D) ARG 217 – GLU 41 salt-bridge formation projected into the energy landscape. The presence of this salt-bridge clearly increases while going from inactive conformation to active-like conformation. But the number is not zero in the inactive conformation like the previous one. What the diagram says is that the salt-bridge isn’t stable in state 2, i.e. the inactive conformation, while in the state 1, i.e. active-like conformation, the salt-bridge is very stable. (E) A representative structure from state 1 where ARG 217 and GLU 41 are situated closely to form a salt-bridge. (F) A representative structure from state 2 where ARG 217 and GLU 41 are situated far apart to form any contact.

Similarly, during the visual inspection using ProteinTools, a strong hydrogen bond network was observed between ARG 217 of the L11-12 region and ASN 37 and GLU41 of the H3 region in active like conformation (Fig S5) which is absent in the inactive conformation. These interactions seem to provide the energetic drive to make the transition from inactive to active-like conformation. These salt-bridges also provide a molecular understanding of the co-related motion between L11-12 and H3 region that was seen from our earlier Dynamical cross-correlation analysis (Fig 1F and 1G).

Interestingly when we checked the trajectories of our active conformation, we didn’t see the presence of these salt-bridges at all. The probability density of the C-alpha distances between GLU 222 and ARG 42 residues, and ARG 217 and GLU 41 residues were plotted for the MD trajectories of both active and inactive conformations (Fig 6 A, B). The range of C-alpha distances is much broader for the inactive conformation compared to the active conformation because of the higher fluctuations of the corresponding residues in the inactive state. It can also be seen that for both sets of C-alpha distances, the peaks for the active state situated in the right-hand side of the 10 Å value signifying the absence of these salt-bridges. To check whether these data are in line with our accelerated MD simulations, we have plotted the similar quantities for our aMD trajectories, and they recapitulate the conventional MD results (Fig S7). In the case of inactive conformation though, the peaks can be seen at much lower values. The free energy surfaces with fraction of native contacts and the distance between the oxygen atom of acidic sidechain and the nitrogen atom of the basic sidechain as the reaction co-ordinates were plotted (Fig S6). In this case we have selected specific oxygen and nitrogen atoms which were seen to be involved in the formation of salt-bridges when visualized in ProteinTools. It can be seen that with the increase in the fraction of native contacts, the distance decreases. But it needs to be mentioned here that the fraction of native, highest being around 0.75, doesn’t really reach the level of the active conformation. Thus, in ideal situations, we need more sampling to see the complete conformational transition. But it still establishes the fact that the salt-bridges between these residues were not formed during the simulations of the active conformation since the minimum distance observed between these atoms there is around 6.5Å, while in the case of the simulations of the inactive conformation we do observe these salt-bridges. If we consider the active conformation as our native structure, which we have discussed previously, these salt-bridges can be termed as non-native salt-bridges. Combining these results with our previous set of results, it can be said that the non-native salt-bridges that were formed near the loop region of the H12 modulate the inactive to active conformational transition in PR.

**Fig 6:**
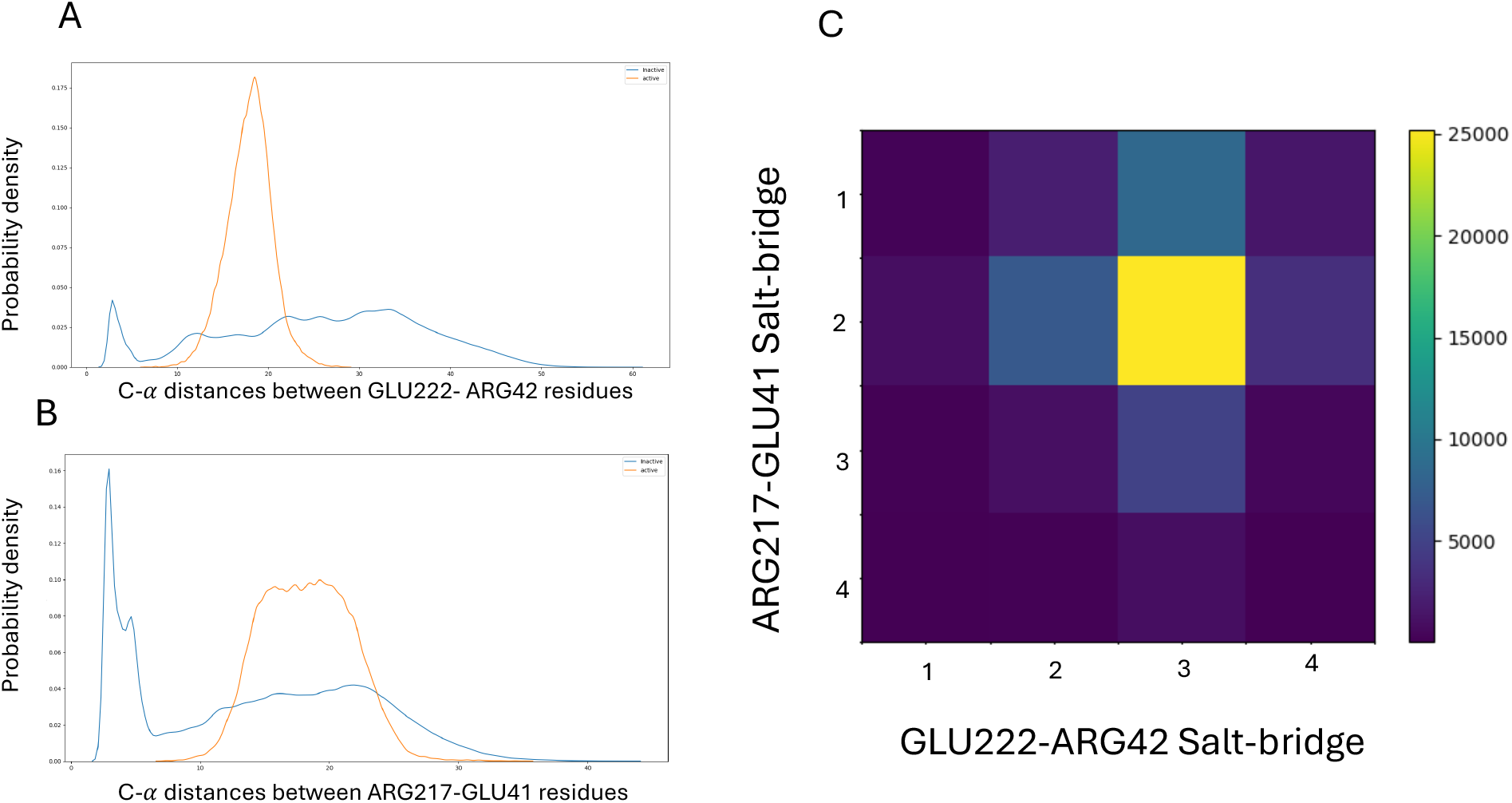
GLU 222 – ARG 42 and ARG 217 – GLU 41 salt-bridges are not found in the simulations of the active conformation. The active conformation is considered here as the native conformation. Thus, these salt-bridges are termed as non-native salt-bridge throughout this paper. (A) The probability density of the C-*α* distance between the residues GLU222 and ARG42 for the two sets of simulations. While for the inactive conformation, there is a peak that lies for a very small value of C-*α* distance, where the salt-bridge can be formed, for the inactive conformation the distance is always too high to form any contacts. (B) The probability density of the C-*α* distances between the residues ARG217 and GLU41. This distance also follows the previous trend. Thus, both these salt-bridges are absent in the simulations of the active conformation. (C) and (D) A graphical representation of all the possible contacts formed by GLU222 and ARG42, and ARG217 and GLU41 residues. In the matrix representation, the rows represent the number of contacts, from 1 to 4, formed between the ARG 217 and GLU 41 residues and the columns represent the number of contacts, from 1 to 4, formed by GLU222 and ARG42 residues. And the color bar represents the number of frames in our trajectories. Most of the frames have both types of contacts present.

Another question that we ask is whether these two salt-bridges are formed simultaneously or successively. We plotted the number of contacts formed by GLU222 and ARG42, and ARG217 and GLU41 residues in each frame in a matrix format in our simulations of the inactive conformation. The greatest number of frames in our combined trajectory has zero values for both these contacts which is understandable since our MD simulations sample the conformational space of the inactive conformation more when the initial conformation is the inactive one (Fig S7). We have also seen regions where ARG217-GLU41 has contacts but not GLU222-ARG42. This reiterates what we have seen from our energy landscape plot, where we saw the formation of GLU217-ARG41 in state 1, i.e. the inactive conformation, but GLU222-ARG42 contacts were absent in this region. Apart from these regions, it can be seen that the dominant region has the presence of both these salt-bridges (Fig 6C). It shows that these two salt-bridges formed simultaneously in state 2, i.e. the active-like conformation, in the energy landscape. Together they seem to provide the energetic advantage to the near-active conformation and thus modulate the conformational transition.

## Discussion

In this paper, we have done a rigorous study of the dynamics of the progesterone receptor when bound to partial agonist Asoprisnil. Using many replicates, we have shown that the inactive conformation shows greater flexibility and greater structural instability in the H12, and L11-12 regions compared to the active conformation, recapitulating our earlier accelerated MD study. The active conformation has a lower SASA value compared to the inactive conformation. A SASA plot over time for one of the trajectories of the inactive complex that shows how the SASA value decreases with time (Fig-S8). The decrease in the SASA value in the H12 and L11-12 regions is indicative of structural stability of the protein.

Because of the large number of replicates, we were able to go beyond simply observing the conformational transition from inactive to active-like conformation and were able to apply the Markov state modelling to study the kinetics of the transition. At a temperature of 310K, this transition happens at a microsecond time scale. Not being able to see a transition into inactive state when started from the active conformation reestablisheses the fact that active conformation is energetically more stable than inactive conformation which we have already shown using umbrella sampling ^14^. The Markov model also helps us to understand the molecular forces that facilitate this transition from the inactive to active-like transition to happen. We have found several interactions like salt-bridges and hydrogen bonding that were shown to be formed by the residues specifically on the L11-12 regions with H3 region that are not present in the inactive structure but starts forming when transition into near-active like conformation. These interactions seem to provide the energetic advantage necessary for the formation of active-like conformation and facilitate the conformational transition.

One of the most significant findings from this study is that the contributing interactions in this transition are non-native, i.e. these salt-bridges are not found to be present in the simulations of the active or native conformation. As we have shown that the active conformation represents the more stable conformation, we termed it as the native conformation and thus the salt-bridges formed are the non-native ones. This study shows how non-native salt-bridges modulate the inactive to active conformational transitions. The importance of non-native contacts in the protein folding is a much-debated one^16–19^. Although our study is not meant to discuss protein folding in general term, it might be influential in dissecting the role of non-native contacts in the bigger problem of protein folding, and not just limited to the question of conformational transition in nuclear receptor.

Considering the importance of Progesterone receptor as a potential drug target for diseases like breast cancer, this study can be used to assist both drug development and protein engineering research. Molecular knowledge can be used to mutate certain residues to either activate or deactivate the receptor, based on the requirements.

Based on all the findings in this study, we have come up with a thermodynamic toy model that might explain this inactive to active conformational transition at the molecular level (Fig 7). It should be emphasized here, as we have discussed previously, our sampling isn’t enough to witness a complete conformational transition process from inactive to active state and hence we combine all our different findings so far to build this simplified heuristic model. In the inactive state of the protein, the atoms of the respective residues are situated far apart. The formation of non-native interactions, salt-bridges and hydrogen bonding, stabilize the structure into an active-like intermediate state which then transitions into the active or native state. During this final transition the interactions are disrupted, and the conformation is stabilized by other factors, for e.g. other interactions and possibly different entropic effects as well. In such a rugged energy surface, the non-native interactions play as a mediator in the conformational transition, providing necessary energetic advantages to the intermediate structures.

**Fig 7:**
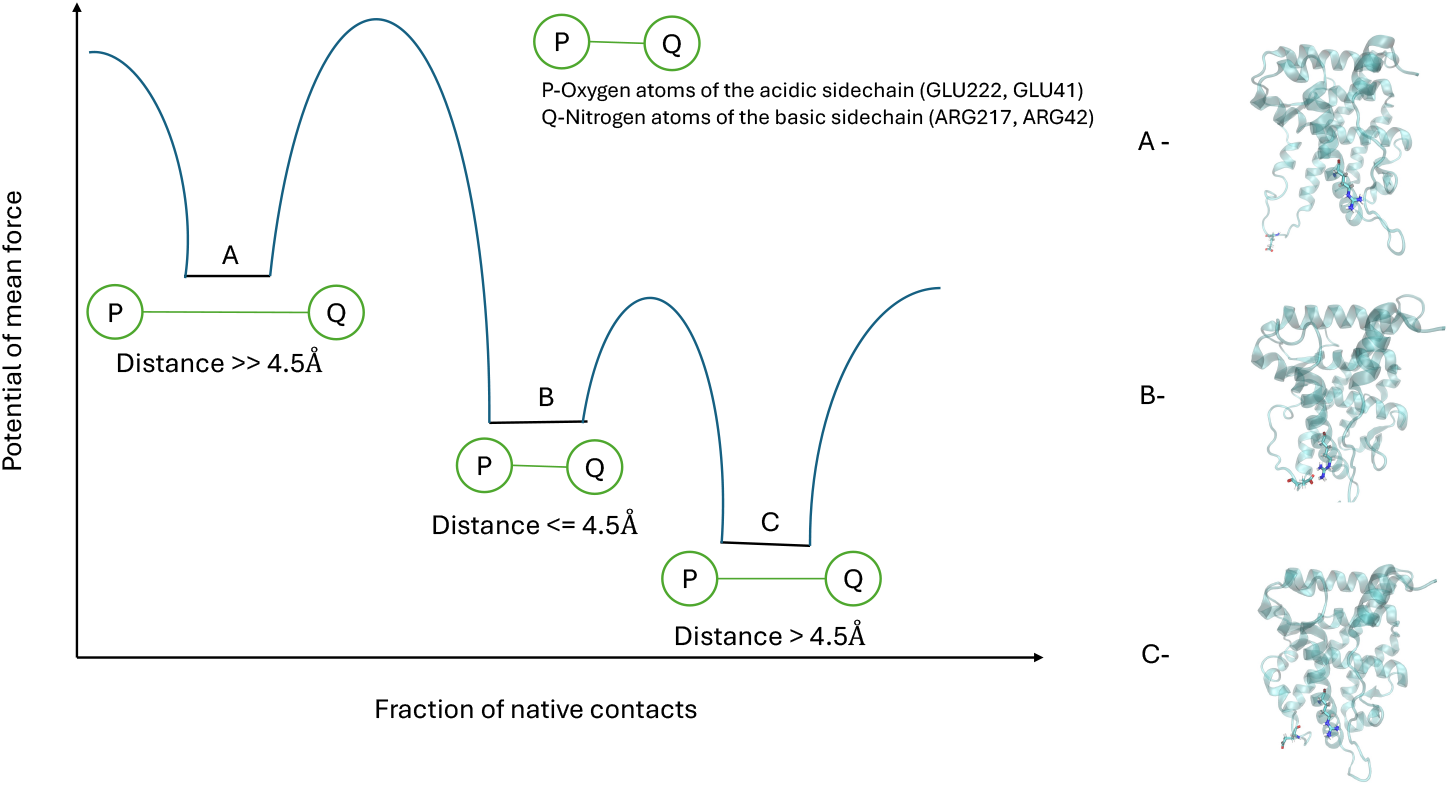
A thermodynamic toy model to explain the inactive to active conformational transition in Progesterone receptor. Inactive conformation, active-like conformation and active or native conformation represent three states in the free energy surface. P-Q represents the distance between the specific oxygen atom of the acidic side chain of GLU222 or GLU41 and specific nitrogen atom of the basic side chain of ARG217 or ARG42.

It’s important to mention here that this study was done for the PR ligand-binding domain only. A more detailed study is needed for the full-length PR. The lack of crystal structure for a full-length PR poses a great difficulty in doing such a detailed study. A full-length PR not only has structured regions, but it also has unstructured hinge and N-terminal domains. Although, in recent times different machine-learning based tools like AlphaFold^30^ has made significant progress in the direction of protein structure prediction for structured regions, it’s still not adequate in predicting the unstructured regions^31^ as IDR often have an ensemble of structures rather than a single structure. Also, PR is functional as a dimeric form and an obvious future direction is to do a detailed study of the dynamics of the PR-dimer in the presence of the DNA response element.

## SUPPLEMENTARY INFORMATIONS

**Fig S1:**
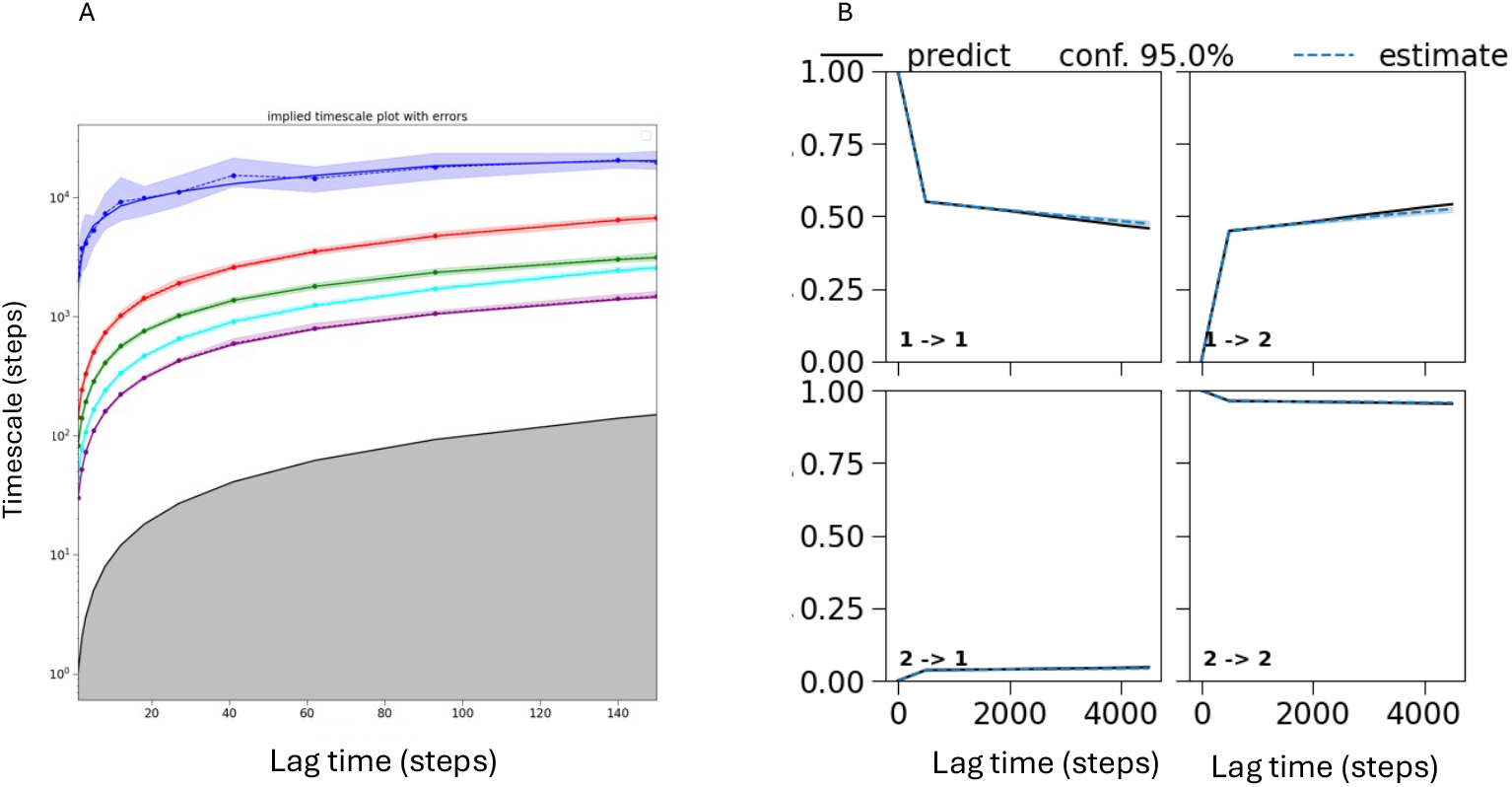
The statistical validation for the MSM of the MD simulations of the inactive conformation. A) Implied timescale of the simulated system. B) Chapman-Kolmogorov test of the system.

**Fig S2:**
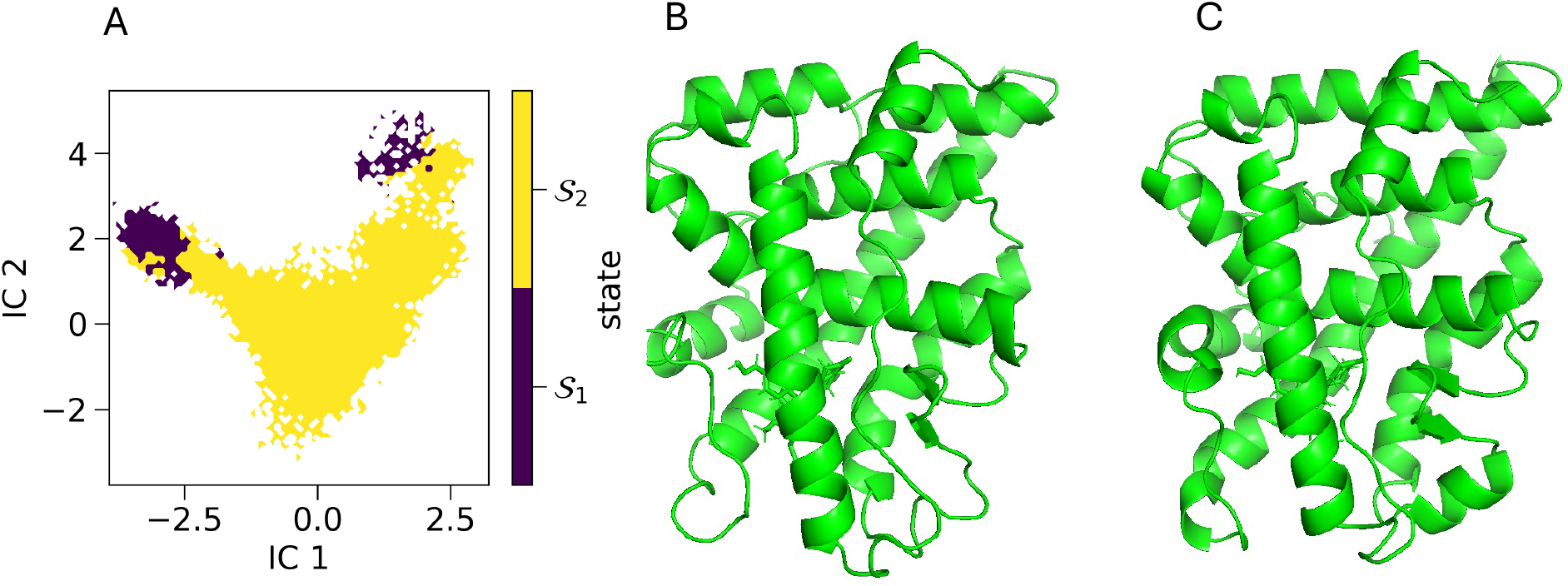
Markov state modeling of PR-active state. (A) TICA plot with two states. (B) and (C) The representative structures from the two states.

**Fig S3:**
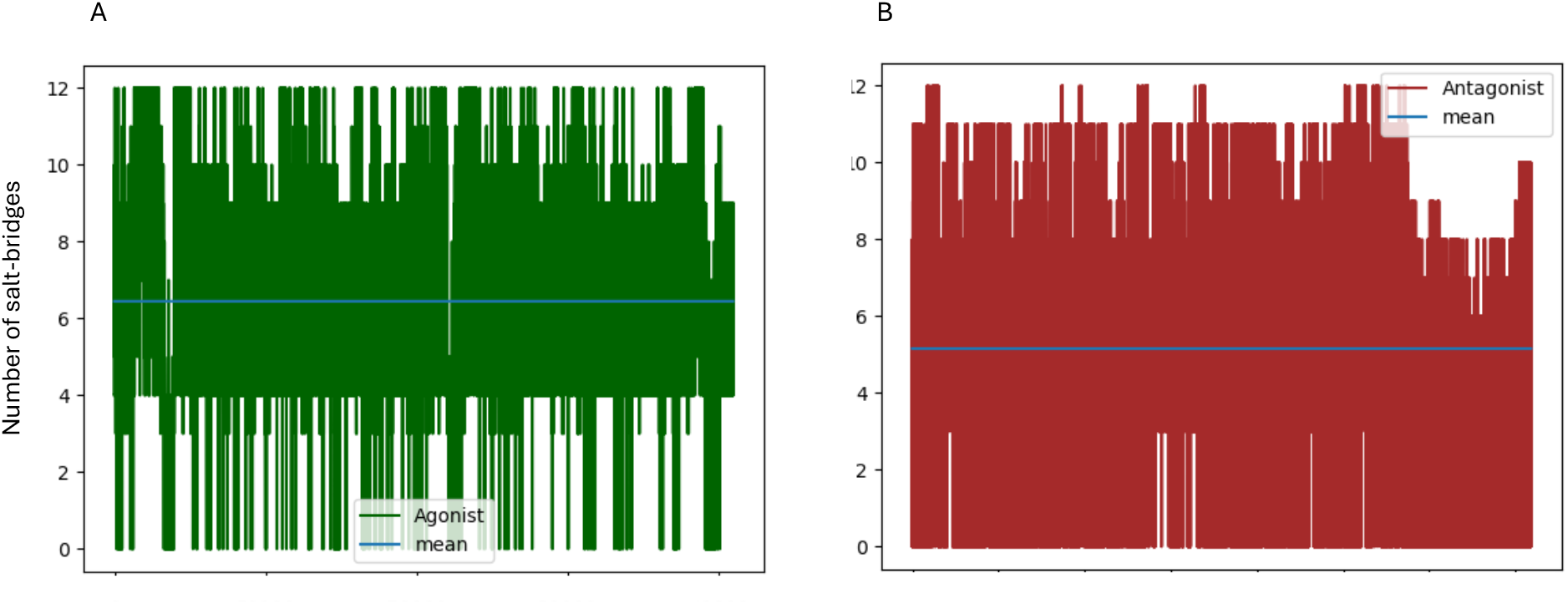
Overall Salt-bridges formed by the H12 region for active (A) and inactive conformation (B). Here in the plots the salt-bridges were calculated for all the trajectories combined. The mean value for the active conformation (6.6) is higher than for the inactive conformation (5.2).

**Fig S4:**
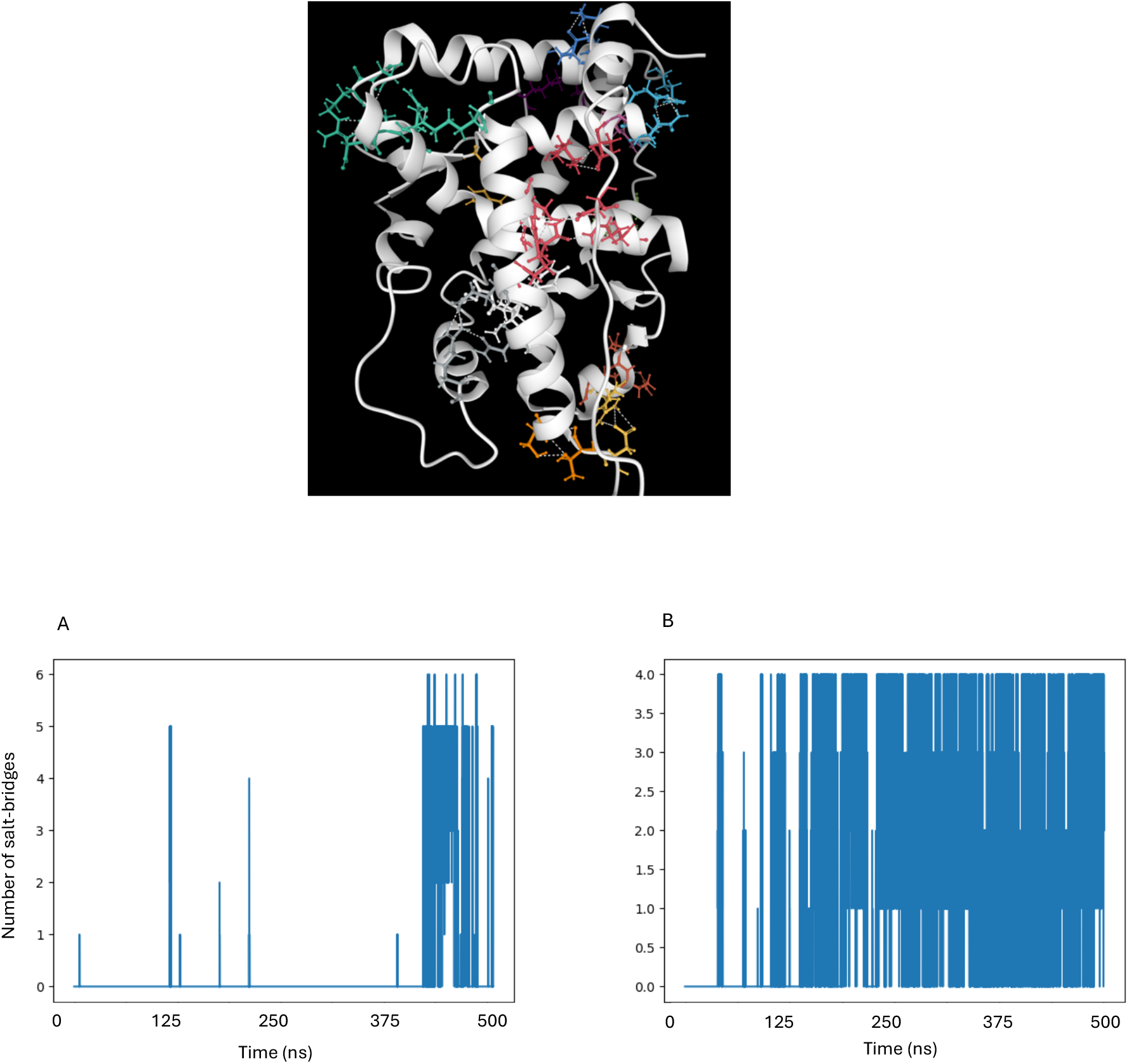
Salt-bridges formation with time for a trajectory of the inactive conformation that shows conformational transition from inactive to active-like conformation. (A) Salt-bridge formation between GLU 222 and ARG 42 increases with time. (B) Salt-bridge formation between ARG 217 and GLU 41 increases with time.

**Fig S5:** The hydrogen bond network formed between ARG 217 of the L11-12 region and ASN 37 and GLU41 of the H3 region in active like conformation

**Fig S6:**
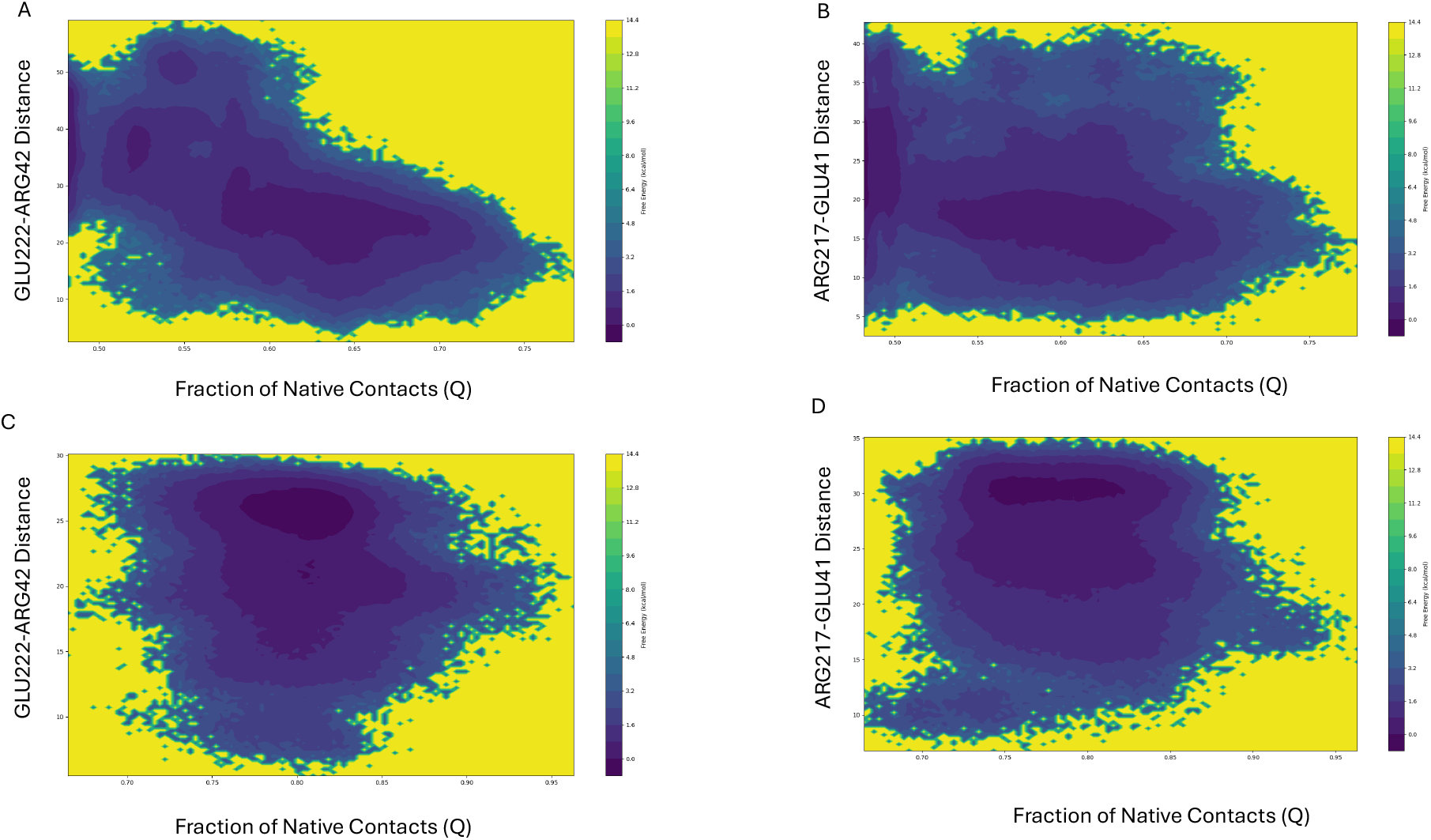
The free energy surfaces with fraction of native contacts and the distance between the oxygen atom of acidic sidechain and the nitrogen atom of the basic sidechain as the reaction co-ordinates were. (A), (B) The 2-D free energy surface for the inactive conformation. (C) and (D) The 2-D free energy surface for the active conformation.

**Fig S7:**
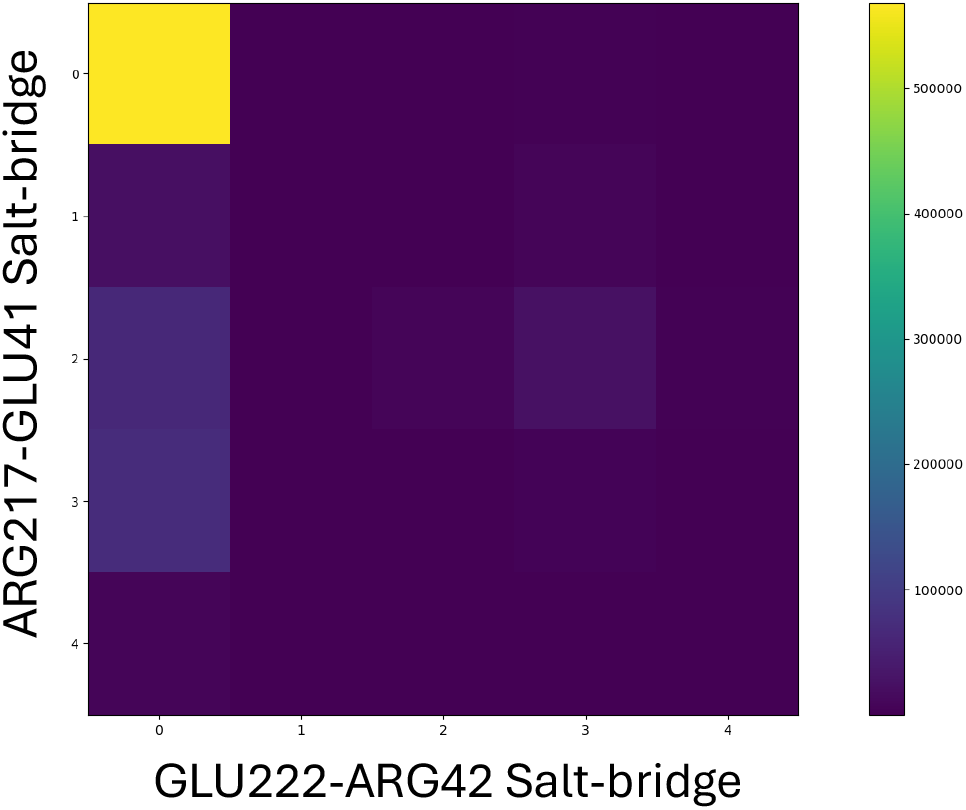
the number of contacts formed by GLU222 and ARG42, and ARG217 and GLU41 residues in each frame in a matrix format

**Fig S7:**
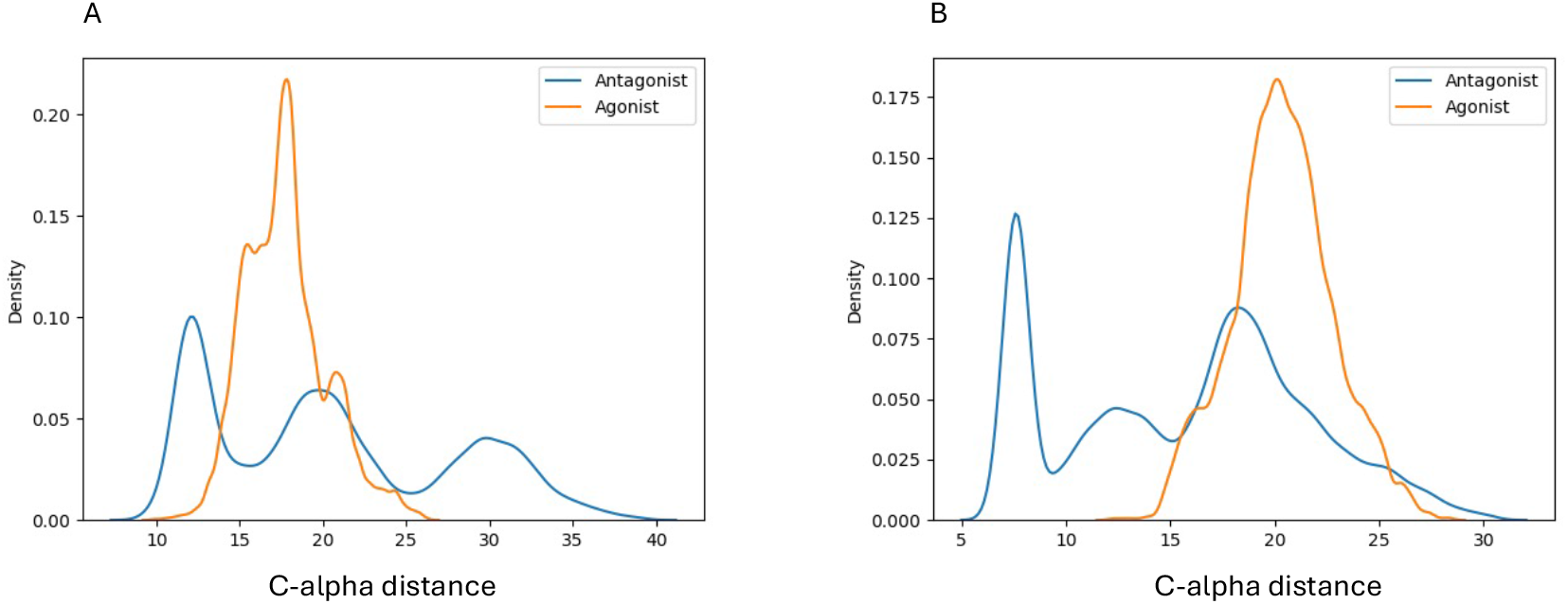
Probability distribution of the C-alpha distances were plotted for accelerated MD simulations. (A) Probability distribution of the *C*_*α*_ distance between GLU 222 and ARG 42 (B) Probability distribution of the *C*_*α*_ distance between ARG 217 and GLU 41.

**Fig S8:**
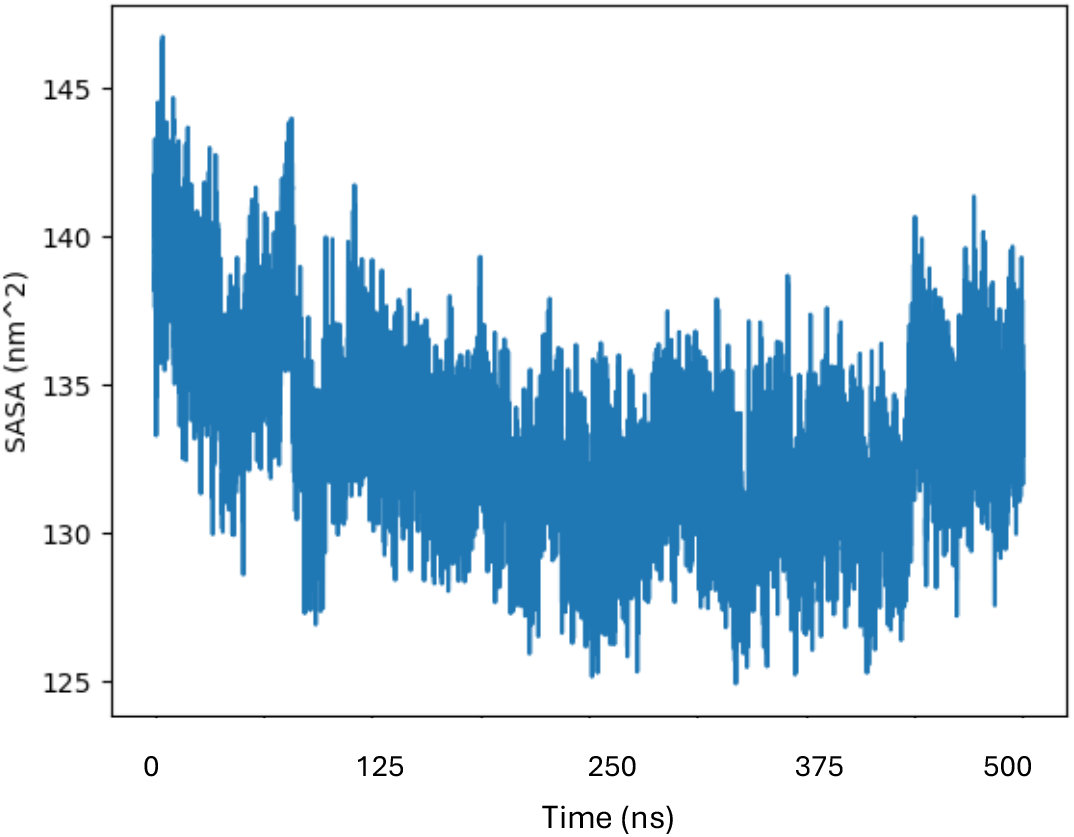
Solvent accessible surface area plot for a trajectory of the inactive conformation that shows the conformational transition from the inactive to active-like conformation.

